# Mathematical Modeling of Bottleneck Transmissions of RNA Virus Infecting a Homogeneous Host Population

**DOI:** 10.1101/2022.08.30.505912

**Authors:** Taimá Naomi Furuyama, Luiz Mario Ramos Janini, Isabel Maria Vicente Guedes de Carvalho, Fernando Martins Antoneli

## Abstract

There is no consensus about when a potential viral infection event presents greater risk of a successful transmission. Some authors suggest that late infection stages present higher risk of transmission. Others suggest that the early infection stages play a most relevant role in transmission events. However, studies considering the fitness or mutational effects on the viral particles over transmission events are lacking. We propose to approach this question through a two-level mathematical model based on RNA viral population dynamics. The first level of the model represents the intra-host viral population dynamics and the second level of the model represents the host-to-host dynamics of transmission events. The intra-host dynamics model uses the fitness of viral particles as means to track the presence of ‘highly infective’ particles during transmission bottlenecks. More specifically, the intra-host dynamics is described by a stochastic quasispecies, based on a multivariate branching process. The host-to-host dynamics of transmission events is emulated by a ‘putative’ transmission tree with ‘host zero’ at the root and a fixed number of branches emanating from each internal node. A ‘Monte Carlo’ strategy was adopted to explore the tree by sampling random walks along transmission chains along the tree. Viral infections of a single host and several transmission events among hosts were simulated in early and late infection stages scenarios. The results show that the early infection stages may represent a key factor in the viral pandemic. Over the evolution of the viral population within each host the mean fitness decreases due to occurrence of mutations (most of them causing deleterious effects). Despite the small opportunity interval, transmissions that occur in early stages could probably infect new hosts at a higher rate than in late stages. It was observed that a very early transmission scenario could reach a transmission chain 20 times longer than a very late transmission scenario. This indicates that the quality of the viral particles is a relevant factor for transmission events.

## 1. INTRODUCTION

According to the UNAIDS Global AIDS Update 2022 report the HIV-1 new infections rate indicates that over 2021, each day, more than 4 thousand people were newly infected. There are about 40 million people living with HIV-1 (human immunodeficiency virus), globally. It is estimated that 85% of people living with HIV-1 knew their status, 75% were under antiretroviral therapy and more than 65% presented a suppressed viral load [1]. To control and decrease the number of newly infected individuals is still a great challenge for several reasons, ranging from economy, social discrimination, cultural behavior for example, and now the SARS-CoV-2 pandemic as well [1–5]. Brazil is at the forefront in Latin America to provide PrEP (pre-exposure prophylaxis) free of charge via a public healthcare system to people included in risk groups [2].

HIV-1 transmission through sexual intercourse represents the most common transmission mode [6–9]. To promote an effective transmission of HIV-1 from one individual to another, viral particles must overcome several physical and chemical barriers imposed by the host, like physical mucosal barriers and secretions, target cell availability, immune response activation, inflammation in exposed tissue, and altered mucosal microbiota [9–12]. Distinct transmission routes represent many different barriers, that is, they compound distinguished transmission bottlenecks between donor and recipient hosts. The mechanisms behind these possible processes are not completely understood [8, 12, 13]. However, it is known that there are specific viral particles responsible for the transmission and establishment of the HIV-1 infection, called the transmitted founder (TF) particles [9, 12, 14].

About 80% of HIV-1 transmissions occur after mucosae exposure (mainly via sexual intercourse) and 60% to 80% of these, the transmission bottleneck allows the establishment of one to five transmitted founder viruses [9, 14–17]. Several viral variants may be present in the genital tract, but only a few particles can be detected in the bloodstream in early stages of infection [12, 14, 18, 19]. An *in vivo* analysis of the exact transmission moment would be unworkable, thus an *in silico* model could be a valuable approach to the study of this problem. The purpose of this paper is to show through mathematical modeling and computational simulations how distinct scenarios of early and late transmission events affect the replicative dynamics of a viral population.

A RNA viral population within a host can be modeled by a dynamical system, called a quasispecies, which contains viral particles with different replicative fitness varying closely to one or more best adapted sequences [9, 20–27]. According to the host’s immunological characteristics, the viral particles may suffer the consequences of the occurrence of mutations, in the form of differential fitness effects. Considering that deleterious and neutral mutations are more likely to occur [28–30], a viral population under high mutation rates may undergo losses in its average replicative capability. The TF particles have features that make them different from the chronic consensus (CC) particles [8–10, 14, 17, 20]. This suggests that at the initial moment of infection the TF articles play a fundamental role in the establishment of the viral quasispecies. The descendants of an early TF particle may lose the qualities of their parentals, but are still able to maintain the survival of the population in a lower replicative level, adjusted to the adaptive landscape shaped by the host’s immune system.

Assuming high mutation rates and quasispecies dynamics, some insights may be gained. If transmission events occur consecutively at short time intervals, that is, with few replication cycles succeeding transmission events, the number of mutations accumulated is expected to be low. Hence, in these early infection events, the population is likely to preserve the characteristics of the TF particles throughout the transmission events inducing more severe conditions.

On the other hand, if the interval between the occurrence of transmission events is very long, the viral population undergoes a high number of replication cycles. Hence, in these late infection events, the viral population is expected to accumulate a high number of mutations (mainly deleterious). In this case, the viral population is likely to lose the characteristics of the TF particles, retaining only mutated sequences that are similar to the TF particles. Therefore, new infections occurring at late infection events possibly transmit only chronic consensus particles and may not even induce an infectious condition on the new host.

Nevertheless, in a real host population these events of transmission can occur at any point in time relative to the intra-host viral population dynamics. The main question we want to investigate is whether the long term effect of higher occurrence of early infection events can be detected at the level of host population.

We propose to approach this question through a two-level mathematical model, a model for the intra-host viral population dynamics and a model for the host-to-host dynamics of transmission events. The intra-host viral population dynamics is described by a stochastic quasispecies model, based on a multivariate branching process introduced and analyzed in [31–33], including its computational implementation. The host-to-host dynamics of transmission events is emulated by a ‘putative’ transmission tree with the ‘host zero’ (H0) at the root and a fixed number of branches emanating from each internal node. The transmission events occur along this ‘putative’ transmission tree from starting at the root and flowing through the internal branches. At each internal branch a random infection occurs, causing a population bottleneck, since only a small number of viral particles are transmitted to the newly infected host. In order to computationally implement the transmission tree part of the model, without running out of computer memory, we adopt a Monte Carlo Markov Chain strategy of exploring the tree by sampling random walks along chains of consecutive branches of the tree.

## 2. MATERIALS AND METHODS

### 2.1. The ENV and the ENV-TF programs

The program described in this paper, called ENVELOPE-TF (ENV-TF), was based on the ENVELOPE (ENV) program (EvolutioN of Virus populations modELed by stOchastic ProcEss). ENV was developed as a computational platform capable of simulating the stochastic dynamics of RNA virus populations, also called quasispecies, in a single host. The ENV program simulates a mathematical model for viral evolution based on the theory of multivariate branching processes and considers only phenotypic properties of a viral population, such as, the distribution of fitness effects and the replicative capability of each viral particle [31–33].

The program ENV-TF uses the same viral populations dynamics of ENV for each host, yet ENV-TF promotes transmission events from one host to another, allowing the phenotypic properties of certain states to be transferred with the viral particles and each state of a limited population of viral particles would represent the viral bottleneck, initializing a whole new viral population within a new host. The program ENV-TF was developed in Python (v3.6) with the Spyder® IDE from Anaconda distribution (v3). The data generated by the program can be exported as an “xlsx” file for further analysis.

The mathematical model underlying the program ENV-TF has two levels: (i) the intra-host viral population dynamics (the same as in the ENV program) and (ii) the host-to-host dynamics of transmission events.

The intra-host viral population dynamics is governed by a multivariate branching process, called the Phenotypic Model, see [31–33]. In this model each viral particle belongs to a replicative class (R), which determines its fitness as the progeny size of the particle. The progeny of all particles is subjected to the same mutational load which can promote deleterious (d), beneficial (b) or neutral (n) effects. These fitness effects affect the replicative class of the progeny particles by decreasing (deleterious), increasing (beneficial) or maintaining (neutral) its value in relation to the replicative class of their parental particle.

For example, a viral particle with replicative class R=3 generates a progeny of size 3. However, these 3 particles do not necessarily have the same replicative class as their parent particle. Each one of them should undergo the effect of mutations according to the probabilities of occurrence of deleterious, beneficial or neutral effects of the model. For sake of simplicity, assume that the three fitness effects have the same probability to occur, that is, d = b = n = 1/3. Since each one of the 3 particles undergoes a mutational effect, it is expected that all the three mutational effects do occur. One particle would undergo a deleterious effect and have class R=2, another particle would undergo a beneficial effect and have class R=4 and another particle would undergo a neutral effect and have class R=3. Thus, the progeny of the particle with class R=3 in this example consists of 1 particle of R=2, 1 particle of R=3 and 1 particle of R=4. All particles in a given generation go through this process simultaneously, giving rise to the viral particles of the next generation.

The host-to-host dynamics of transmission events is defined in terms of a mechanism of viral transmission from an infected host to a non-infected host. Hence, it must include several hosts and many directions of transmission, readily becoming a complex net of interactions among several hosts. A simplified interaction system among hosts is given by a transmission tree. In such a scheme, one infected host would infect a fixed number M of hosts (at different times), and each one of these M hosts would infect other M hosts and so on, generating the infected population with growth rate given by a geometric progression.

The main problem in implementing a full transmission tree is the exponential growth of the number of hosts that the program must keep track. We solve this problem by sampling the ‘putative’ transmission tree by random walks along linear chains of infection. That is, we start with 1 initial infected host, called H0, that at a certain replication cycle infects another host which has no previous infection, and this last host infects another e so on, forming a linear chain of infected hosts. Each chain is a random walk on the transmission tree since the transmission times are random. Although a single random walk is not representative of all the possibilities of the transmission tree, a sufficiently large set of random walks should be able to explore almost all the possibilities of the transmission tree. Therefore, each complete simulation performed in this paper consists of a large number of random walks which aim to approximate a transmission tree representing a certain transmission scenario (see Fig. 1).

**Figure 1.**
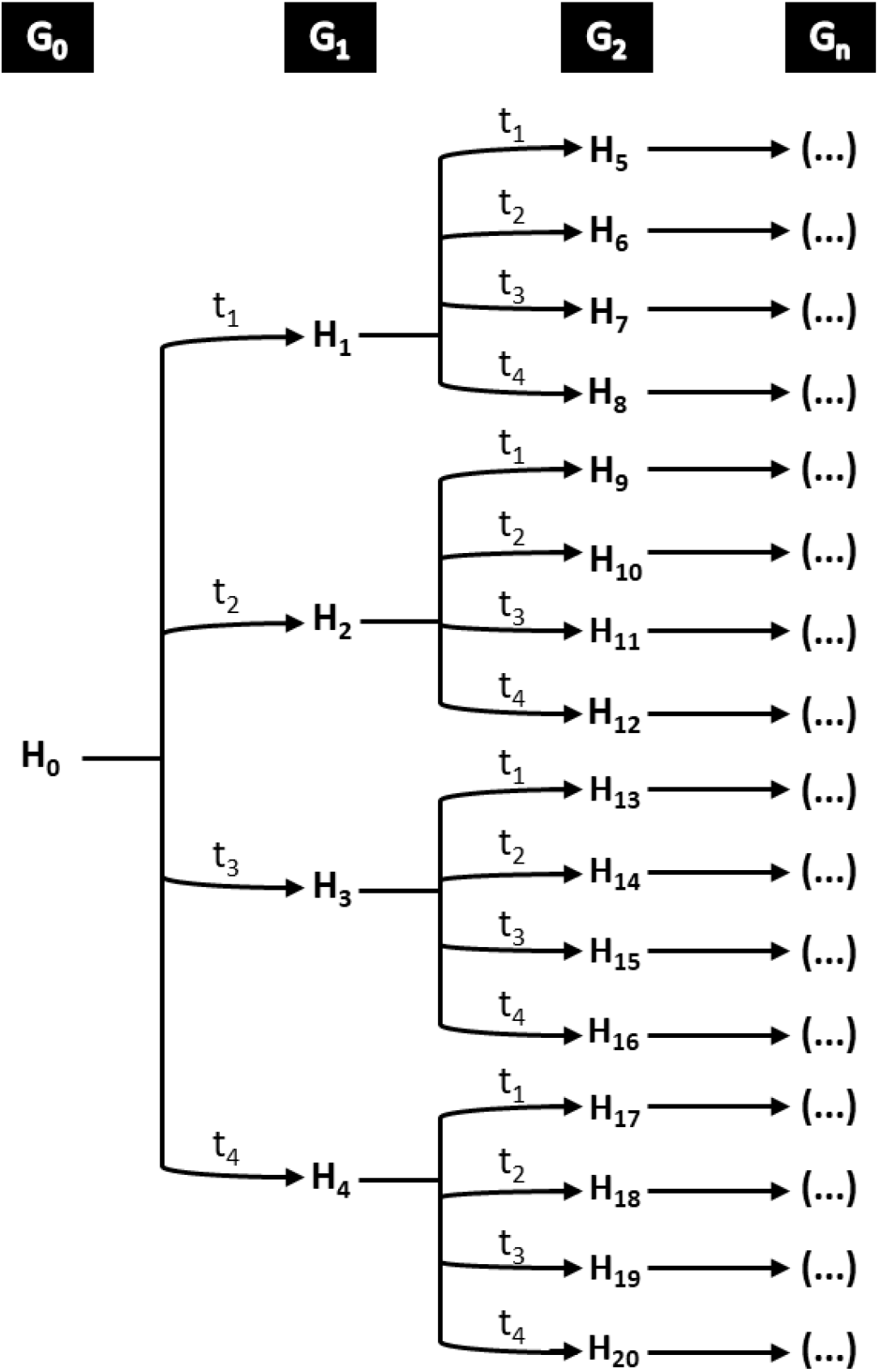
A ‘putative’ transmission tree scheme where the paths represent possible random walks along chains. The program implements this scheme by considering that each simulation represents one random walk given by a single transmission chain. For instance, one simulation of a very early scenario could be from host zero (H0), to H1, to H5 and so on. A late transmission scenery would go from H0, to H3, to H15 and so on. The set of 8,000 simulations intends to cover all the theoretical available possibilities in order to provide an unbiased dataset. Each infected host infects four new hosts in different moments in this theoretical tree. The first four hosts infected form the first generation of hosts; H5 to H20 are the second cluster of infected hosts and form the second generation and so on. Regardless of the moments of transmission, each generation is defined by the numeric order of the transmission event.

The input parameters for the intra-host dynamics is (see [31] for details):

1. Maximum Number of Replication Cycles (Tmax): it is a positive integer greater than one that defines the maximum number of replication cycles in a simulation. A replication cycle is the necessary time interval for a viral particle to be able to give rise to new viral particles, that is, the time interval to give rise to the viral progeny.
2. Maximum Number of Replicative Classes (R): it is the fitness attribute of each viral particle. It is defined as the progenie size produced by a viral particle. The replicative class is represented by R and varies from zero (R=0) to a maximum value of Rmax.
3. Beneficial Mutation Probability (b): it is a real number between 0 and 1. The occurrence of a beneficial effect during the production of a progeny particle increases by 1 the replicative class of the progeny in relation to its parental particle.
4. Deleterious Mutation Probability (d): it is a real number between 0 and 1. The occurrence of a deleterious effect during the production of a progeny particle decreases by 1 the replicative class of the progeny in relation to its parental particle.
5. Neutral Mutation Probability (n): it is a real number between 0 and 1. The occurrence of a neutral effect during the production of a progeny particle does not change the replicative class of the progeny in relation to its parental particle. In the program, the three mutation probabilities (beneficial, deleterious and neutral) satisfy the relation n = 1 – d – b.
6. Maximum Number of Viral Particles per Replication Cycle (Kmax): it is the maximum number for the viral particles allowed at each replication cycle, also known as carrying capacity. When the quantity of viral particles produced in one replication cycle surpasses this limit the program excludes the surplus by a random sampling process.
7. Replicative Class of the TF Particles of Host Zero: the replicative class of the Transmitted Founder initial particles of Host Zero.

The main output data generated by the intra-host dynamics is:

A discrete probability distribution supported on the set of replicative classes {0,…,Rmax}, called Replicative Classes Distribution. This probability distribution is defined by computing, for each replicative class R, the probability that a randomly selected particle in the population belongs to the replicative class R. The Malthusian Parameter or Average Replicative Capability, denoted by µ. In the phenotypical model proposed in [31] the Malthusian Parameter µ is defined as the mean value of the Replicative Classes Distribution. In general, the Malthusian Parameter µ of a branching process is defined as the mean progeny size of the viral population. It can be shown that for the phenotypical model proposed in [31], where the Malthusian Parameter is defined above, both definitions are equivalent.

It is a general result of the theory of branching processes that the Malthusian Parameter controls the ultimate fate of the population. If µ > 1 then, with probability 1, the population will grow indefinitely with growth rate µ. In this case, the branching process is called *supercritical*. If µ < 1 then, with probability 1, the population will become extinct, in finite time. In this case the branching process is called *subcritical*. When µ = 1, the branching process is called *critical*. In a critical branching process the extinction of the population is certain with probability 1, but the average time for it to become extinct is infinite.

When there are no beneficial mutations – i.e., there are only deleterious and neutral mutations – the Malthusian parameter µ can be explicitly calculated and is given by µ = (1 – d) Rmax. In particular, in a mutation-free environment, the Malthusian Parameter µ is the maximum replicative capability Rmax.

The input parameters for the host-to-host transmission dynamics is:

1. Transmission Cycle: the user must define at which moments, measured in replication cycles, the transmission events should occur.
2. Transmission Bottleneck Size: is the number of viral particles that will be transmitted from the donor host to a recipient host.
3. Number of Infected Hosts: number of hosts that receive particles through transmission bottleneck.
4. Number of Random Walk Chains: the user defines the quantity of random walk chains the program should run.

### 2.2. Validation ENV-TF program

The description above gives the common features between the programs ENV and ENV-TF and such commonalities were employed to validate the ENV-TF program. In both programs, the distribution in the replicative classes can be explicitly determined in the case where the beneficial probability is zero. It is a binomial distribution BINOMIAL(1 − d,Rmax+1), where n = 1 − d is the neutral probability and Rmax the maximum replicative capability [31]. The viral replication dynamics was evaluated for each program according to the following configuration: 100 simulations of a single host, with 99 cycles each, beneficial probability b = 0.008 and deleterious probability d = 0.8 (by complementarity, neutral probability n = 1.992). The result of these simulations displayed in Fig. 2 shows that both programs were in very good agreement.

**Figure 2.**
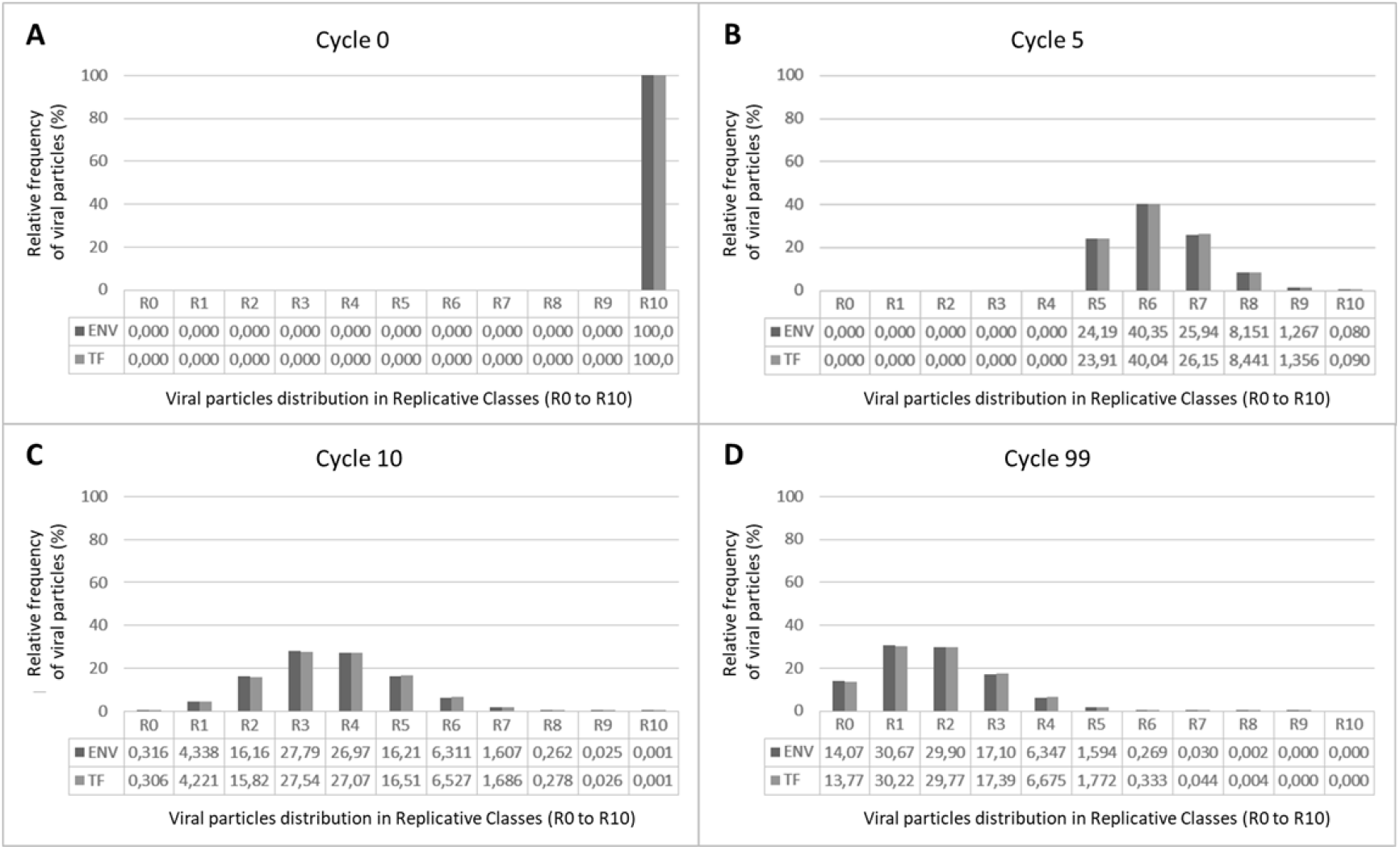
Comparison of the distribution of replicative classes computed by the ENV-TF program (“TF” in clear gray, bars on the right) and the ENV program (“ENV” in dark gray, bars on the left). The computed values are mean values of 100 simulations. Each simulation represents one single host with 99 replication cycles. The probability of occurrence of beneficial mutation was 0.08, of deleterious mutations was 80 and of neutral mutations was 19.92. Each panel shows the viral particles distribution over the replicative classes R (which vary from R0 to R10) in a certain replication cycle (in cycle 0 (A), in cycle 5 (B), in cycle 10 (C) and in cycle 99 (D). The distributions for both programs were in agreement.

As a quantitative validation, the Kullback-Leibler (KL) divergence was computed in order to evaluate how much the replicative classes distributions estimated by the ENV and the ENV-TF programs diverge from each other. If the KL divergence is close to zero, the distributions are considered to be very similar; in an opposite way, if the KL divergence is close to 1, the distributions are considered to be distinct [34]. In each program one simulation was performed, for a single host, with 1,000 replication cycles, 0.0001% of beneficial mutation probability and 50% of deleterious (and 49.9999% of neutral, in a complementary way). The data obtained from cycles 900 to 999 were considered for the estimation of KL divergence. This is enough to make the branching process essentially stationary (actually, it is the distribution of replicative classes that become stationary). The mean value for the KL divergence between the ENV and the ENV-TF replicative classes distributions was −3.25×10^−4^ with a variance of 1.89×10^−6^.

### 2.3. Simulation strategy using the ENV-TF program

Each replication cycle of the ENV-TF program contains the progeny information, such as the number of viral particles and the replicative class of each particle. To follow all transmission events, it is necessary to keep track of the information regarding the replicative classes at each transmission bottleneck.

The input data employed in the intra-host dynamics simulations was:

1. Maximum Number of Replication Cycles (Tmax): we used Tmax = 45 replication cycles per host (cycles 0-44). This number was chosen considering the reference that one replication cycle of HIV-1 takes about 2.6 days [35, 36].
2. Maximum Number of Replicative Classes (R): the default value of Rmax=10 was used.
3. Beneficial Mutation Probability (b): in all simulations this value was fixed at 0.03% during cycles 0 to 8 and increased to 0.08% from cycle 9 onwards.
4. Deleterious Mutation Probability (d): in all simulations this value was fixed at 30% from cycle 0 to 8 and increased to 80% from cycle 9 onwards.
5. Neutral Mutation Probability (n): in all simulations this value was fixed at 69.97% during cycles 0 to 8 and decreased to 19.92% from cycle 9 onwards.
6. Maximum Number of Viral Particles per Replication Cycles (Kmax): the default value of Kmax=1,000,000 was used.
7. Replicative Class of the TF Particles at Host Zero: the default of value R=10 was used.

The input data employed in the host-to-host dynamics simulations was:

1. Transmission Cycle: we used 4 transmission times for each host: cycle 4 (very early), cycle 13 (early), cycle 24 (late) and cycle 44 (very late).
2. Transmission Bottleneck Size: we used an initial inoculum of 5 particles. According to some papers that sought to estimate the effective size of transmission bottlenecks of HIV-1, the number of TF particles is 1-5 [8–10, 14].
3. Number of Infected Hosts: in our simulations we used chains of 101 hosts (H0 to H100) or 501 hosts (H0 to H500).
4. Number of Random Walk Chains: we used 1,000 random walks per simulation for the chains of length 101 and 350 random walks per simulation for the chains of length 501.

From the parameters given in ‘Transmission Cycle’ the transmission tree can be defined through the generation of several transmission chains sampled according to the parameters.

For instance, suppose there is a donor (infected) host A and at a certain moment, say in the 4th replication cycle, a transmission event occurs. The receptor (non-infected) host B receives a small number of the viral particles present in the donor at the 4th replication cycle. Consider that the *transmission inoculum* is of 5 viral particles. The viral particles that compose the transmission inoculum are sampled randomly from the whole viral population in the donor at the 4th replication cycle. The only information that is needed is about their replicative class. If A transmits 3 particles of R=8 and 2 of R=9 then this is the fitness composition of the initial population that starts the infection in B. Continuing this process, at some other moment, B transmits an inoculum of 5 particles to a new host C. Suppose that the transmission from B to C occurs at the replication cycle 40 of B. The viral population in B went through several mutation events, especially deleterious. It is expected that the viral particles that are sampled from B at this replication cycle have lower replicative capability. This process continues forming a chain of transmission events (see Fig. 3).

**Figure 3.**
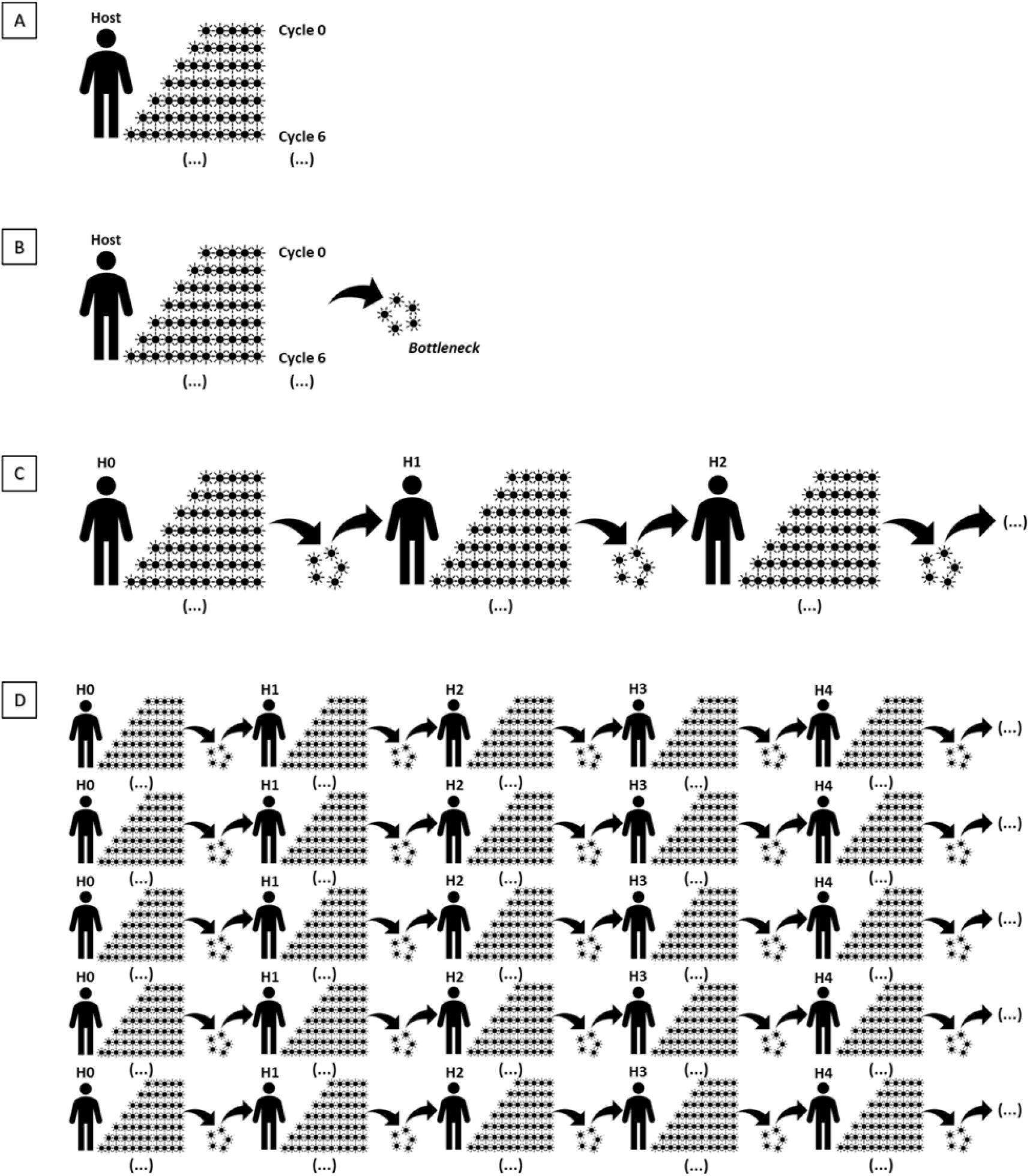
Panel A illustrates a general outline of how the program works for one host only. The initial host starts with a viral population with 5 particles. The image indicates that there are 5 particles at cycle zero, which generates progenie up to replication cycle 6, although the replication process continues until a maximum replication cycle (the ellipsis indicates a continued process). Panel B illustrates the formation of a viral bottleneck, which is given by randomly sampling 5 viral particles from the population at a certain replication cycle. The transmission moments are determined by the user. Panel C illustrates the initial segment of a chain of transmissions, where the initial host (host zero or H0) starts the infection process with five viral particles of class R=10. Over a certain number of replication cycles, the viral population grows, subjected to mutation events. At a certain moment (measured in replication cycles), a bottleneck is formed and a transmission event occurs. An inoculum of 5 viral particles is transmitted to a new host, called H1. The replicative classes of these 5 viral particles are not necessarily R=10, since there were occurrences of mutations during the generation of progenie in H0. The same process now happens with H1 in place of H0 and H2 in place of H1. Panel D illustrates several chains of transmissions representing the random walks on the transmission tree. Each one of these random walks differ (possibly) by the time interval that the transmission events occur.

The transmission chains formed in this way can be interpreted as random walks on a fixed transmission tree, as the one shown in Fig. 1. By generating several such random walks one can explore almost all the possibilities of the transmission tree. The randomness of a transmission chain is given by the choice of time interval at which the transmission occurs. In this paper we consider 4 time intervals (measured in replication cycles): 0-4 (very early), 5-13 (early), 14-24 (late), 25-44 (very late). By defining distinct probabilities for the occurrence of transmission at these time intervals we can generate distinct *transmission scenarios* (TS). These scenarios represent different strategies from exploring the transmission tree.

The total number of hosts considered in the transmission tree can be estimated using the sum of a geometric progression. Equation (1) below gives the total number of infected hosts *a*_*n*_ at the *n*-th generation.

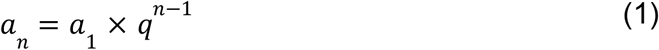

where the common ratio of the geometric progression *q* is the number of infections that each host is able to promote. In this paper, we considered that there was a unique first donor host, the host zero (H0) and so *a*_1_ = 1 in eq. (1).

In a transmission chain, one generation is defined by the occurrence of a transmission event. In the transmission tree, a generation is given by the cluster of hosts infected after the occurrence of a transmission event. For instance, the first generation comprises all hosts infected after the first event of transmission in all transmission chains (see Fig. 1). To calculate the total number of infected hosts in a transmission tree, we compute the sum *S*_*n*_ of the terms of the finite geometric progression, as shown in equation (2) below

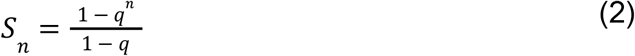

For example, considering that in our simulations each transmission chain had length *n* = 101 and each host infects 4 other hosts, we have that

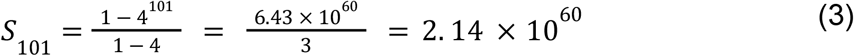

That is, the total number of individuals represented by a transmission tree with *n* = 101 and *q* = 4 is *S*_110_ ≈10^60^.

Of course, the construction of the actual transmission tree is never performed. That is why we call it a ‘putative’ transmission tree. The transmission tree is only ‘realized’ through the exploration of transmission chains during the simulations. To increase the probability of exploring all possible walks along the tree, more than 8,000 simulations were performed.

We tested nine transmission scenarios (named TS1-TS9), each one of them characterized by distinct probability distributions according to the replication cycle interval at which the transmission occurs (see Table 1). We used 5 viral particles with replicative class R=10 (here the replicative classe range from R=0 to R=10) to start the infection in the first host (H0). These 5 initial particles represent the Transmitted Founder (TF) particles.

**Table 1.**
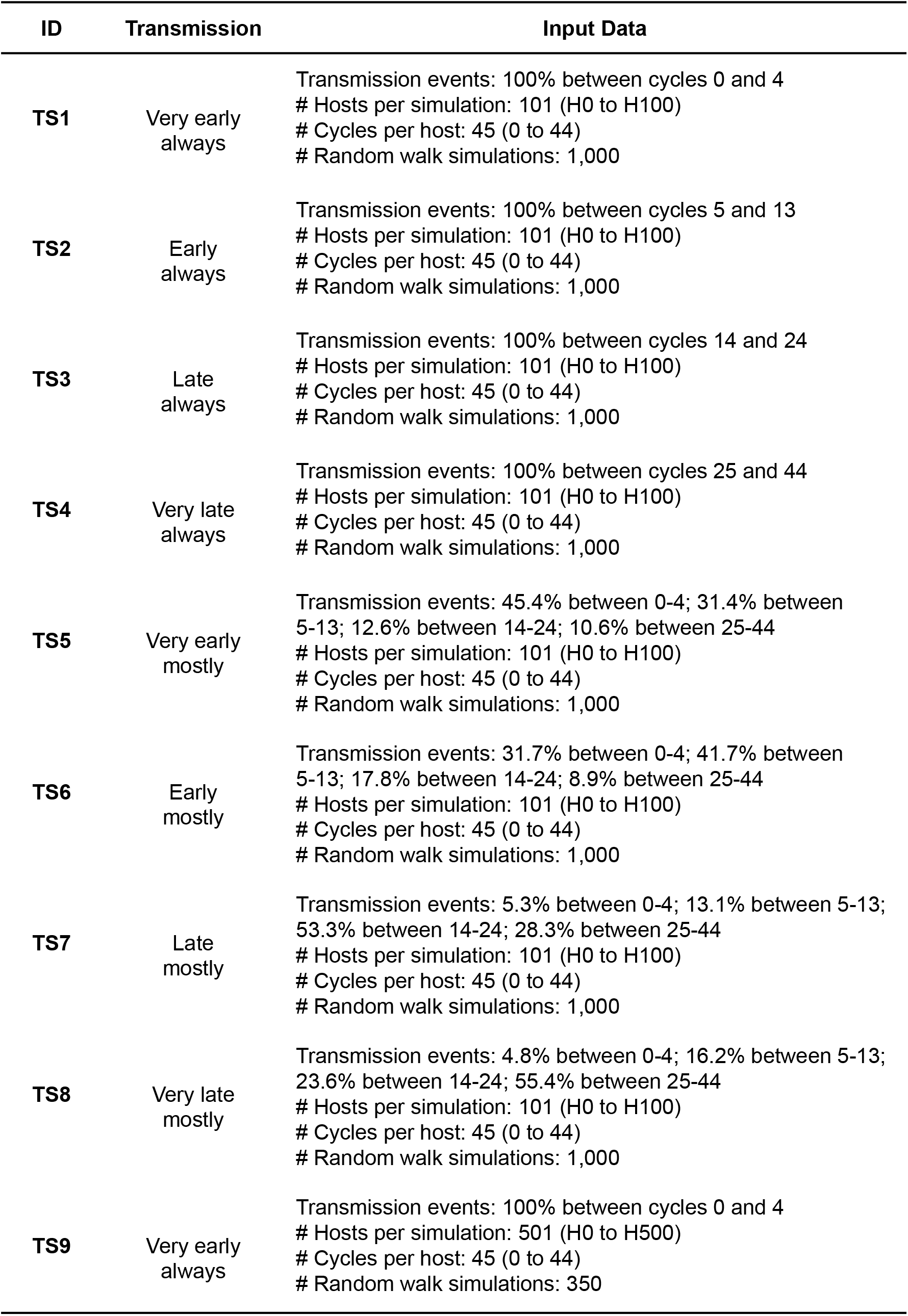
Identification of all Transmission Scenarios (TS) simulated in this paper. Here, the ID column is the name of the Transmission Scenario. For each ID, the Transmission column is the main characteristic of the Transmission Scenario. The column Input Data defines the data that was used in the simulations of each Transmission Scenario. All simulations were executed with the same values for the other parameters available in the program.

The first 4 transmission scenarios (TS) have completely inhomogeneous transmission rates, differing only on the times of transmission. Scenario TS1 has transmissions occurring with probability 1 between cycles 0 and 4. That is, every host of the transmission chain transmitted 5 viral particles between cycles 0 and 4. Scenario TS2 has transmissions occurring with probability 1 between cycles 5 and 13. Scenario TS3 has transmissions occurring with probability 1 between cycles 14 and 24. Scenario TS4 has transmissions occurring with probability 1 between cycles 25 and 44.

The next 4 transmission scenarios allowed for a variation in the probabilities of transmission occurrence among the replication cycles. Scenario TS5 has a probability of 4.54 that transmission occurs between cycles 0-4; a probability of 3.14 that transmission occurs between cycles 5-13; a probability of 1.26 that transmission occurs between cycles 14-24 and a probability of 1.06 that transmission occurs between cycles 25-44. Scenario TS6 has a probability of 3.17 that transmission occurs between cycles 0-4; a probability of 4.17 that transmission occurs between cycles 5-13, a probability of 1.78 that transmission occurs between cycles 14-24 and a probability of 0.89 that transmission occurs between cycles 25-44. Scenario TS7 has a probability of 0.53 that transmission occurs between cycles 0-4; a probability of 1.31 that transmission occurs between cycles 5-13; a probability of 5.33 that transmission occurs between cycles 14-24 and a probability of 2.83 that transmission occurs between cycles 25-44. Finally, scenario TS8 has a probability of 0.48 that transmission occurs between cycles 0-4; a probability of 1.62 that transmission occurs between cycles 5-13; a probability of 2.36 that transmission occurs between cycles 14-24 and a probability of 5.54 that transmission occurs between cycles 25-44. In all of these 8 scenarios the random walk chains contained 101 hosts, from H0 to H100, for each host the number of intra-host replication cycles was 45, from 0 to 44, and 1,000 random walk chains were simulated.

The ninth transmission scenario, TS9, was defined with transmissions occurring with probability 1 between cycles 0-4. The main difference between scenarios TS1 and TS9 is that the former had 101 hosts chains with 1,000 random walks simulated and the latter had 501 hosts chains with 350 random walks simulated. See Results and Discussion for a justification why this last scenario was considered.

### 2.4. Analysis of the data generated by the ENV-TF program

The data collected from all the simulations were analyzed taking into account the parameter values at replication cycle zero and at the last replication cycle (44th cycle). The values at replication cycle zero inform the fitness of the particles at the transmission bottleneck and the last replication cycle informs the fitness of the particles that were susceptible to the mutation probabilities within each host. The data of a simulation generated by ENV-TF can be exported in “xlsx” format. Each sheet of an “xlsx” file corresponds to one host. In order to summarize the data from all simulations, a semi-automated analysis was performed by a Python script generating one single “xlsx” file.

### 2.5. Simulation benchmark and machine configurations

The total time spent on simulations was about 336 hours (during 24 days) in a desktop Intel i5 4690 (4th gen), CPU 3,5 GHz, 8 GB RAM memory, operational system Windows 10 - 64 bits and in a laptop Dell, Intel i5 6200U (6th gen), CPU 2,3GHz, 8GB RAM memory, operational system Windows 10 - 64bits.

## 3. RESULTS

The progenies generated in the program represent only viable or replicative viral particles (it excludes defective particles and does not represent the viral burst size). Genetic drift effects are represented by the bottleneck effects over the infected hosts [17, 37]. While the particles are transmitted, their fitness information is retained in a way that the next infected host starts its viral population with the same number of particles and variable fitness values, according to the fitness obtained from the donor host. The cycle zero from each host reflects the oscillations in the fitness of the bottleneck throughout the transmission events.

The data obtained from the model described in this paper allowed us to analyze the fitness of Transmitted Founder particles according to their Rmax and µ values (see Materials and Methods section). Therefore, it was also possible to verify how many hosts were infected in a transmission chain. The Transmission Scenarios (TS) were divided according to the transmission moment: very early (TS1, TS5 and TS9), early (TS2 and TS6), late (TS3 and TS7) and very late (TS4 and TS8), see Table 1.

Scenario TS1 (very early transmission always) comprises transmission events occurring only between cycles 0 and 4. At each transmission event, a replication cycle in this range was randomly chosen to be sampled for the inoculum. The 1,000 simulations reached 21 hosts (H0 to H20) in the transmission chain and 84 simulations reached 101 hosts (H0 to H100), shown in Fig. 4A (x axis). The Rmax parameter, see Fig. 4A (y axis), shows a steep decrease from H0 to H20, attaining Rmax=5.1 at H20. The mean values of the parameter µ in the first replication cycle (µ_first_) were in agreement with the Rmax mean values. On the other hand, after H20 the parameter µ_first_ decreased gradually till the end, with µ_first_=4.2 at H20 and µ_first_=1.3 at H100 (Fig. 5A). The branching process status (whether µ > 1 or µ *≤* 1 in the last replication cycle) showed a supercritical status until H72. From H73 onwards the status fluctuated from supercritical, to critical and finally subcritical (Fig. 6A).

**Figure 4.**
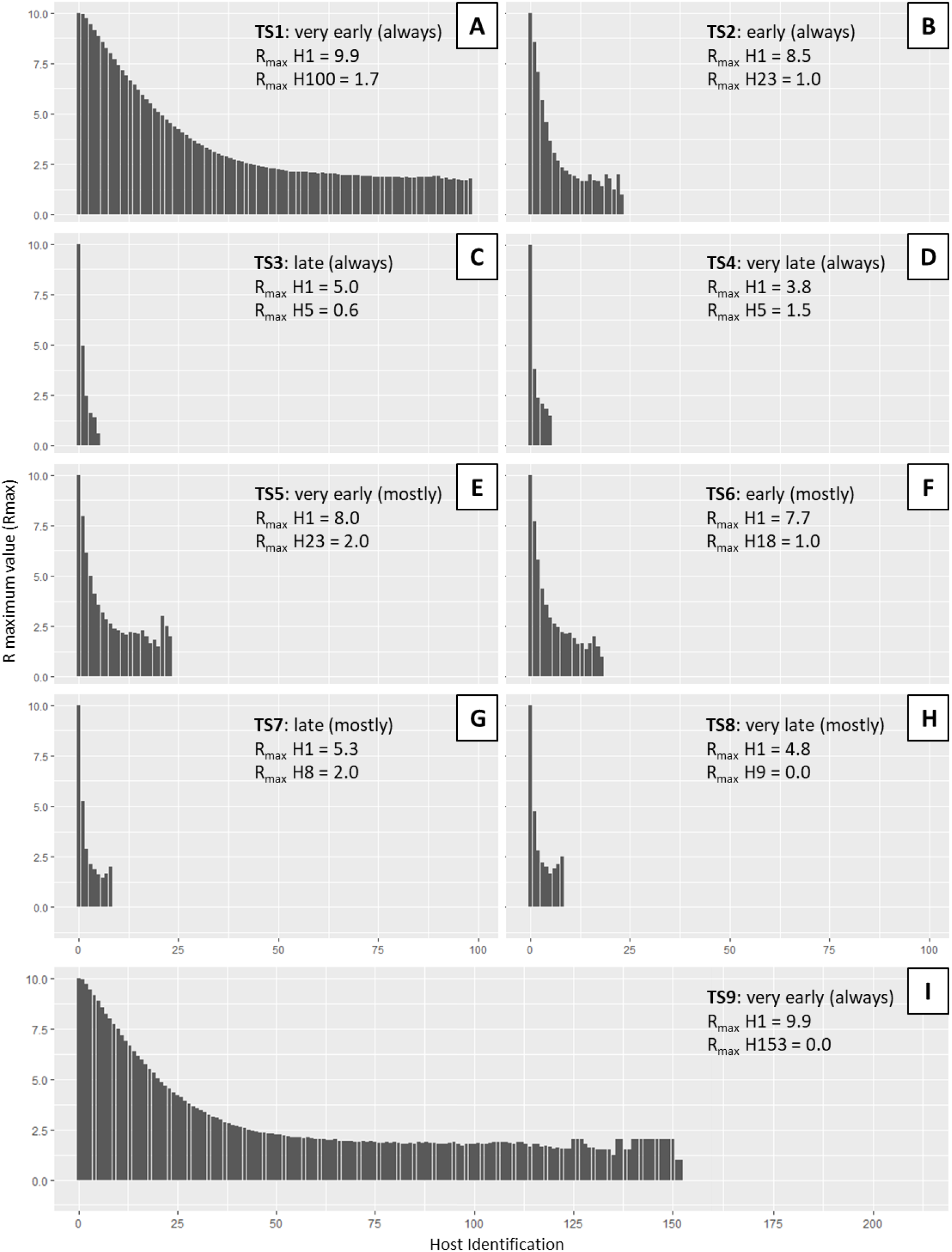
Mean R maximum (Rmax) values of the 1,000 simulations for each scenario, from TS1 to TS9. Each simulation (from TS1 to TS8) was programmed to reach 101 hosts and 44 replication cycles within each host by default in the transmission chain. The TS9 was built with 350 simulations and 44 replication cycles for 501 hosts. The y axis, the “R maximum value” label, shows the values of the replicative classes that ranges from zero to 10 (R0 to R10) and the x axis, “Host Identification”, indicates the amount of infected hosts in the simulations, ranging from the host zero (H0) to the host 100 (H100) and H153. The first host, H0, is the one that harbors the five higher fitness TF particles that replicate into a viral population and initiate the transmission chain. In the very early and early scenarios (TS1, TS2, TS5, TS6 and TS9) the following hosts still receive high fitness viral particles within a short transmission time. In the opposite way, for the late and very late transmission events (TS3, TS4, TS7 and TS8), the Rmax value suffers an abrupt decline in the second host already (H1). This indicates that the viral population within the donor host underwent several replication cycles and consequently several mutation events. For this reason, the TF features were lost and the high replication classes were diminished, reaching a shorter transmission chain. Each panel shows the Rmax value for the H1 and the last host infected in the summarized data text.

**Figure 5.**
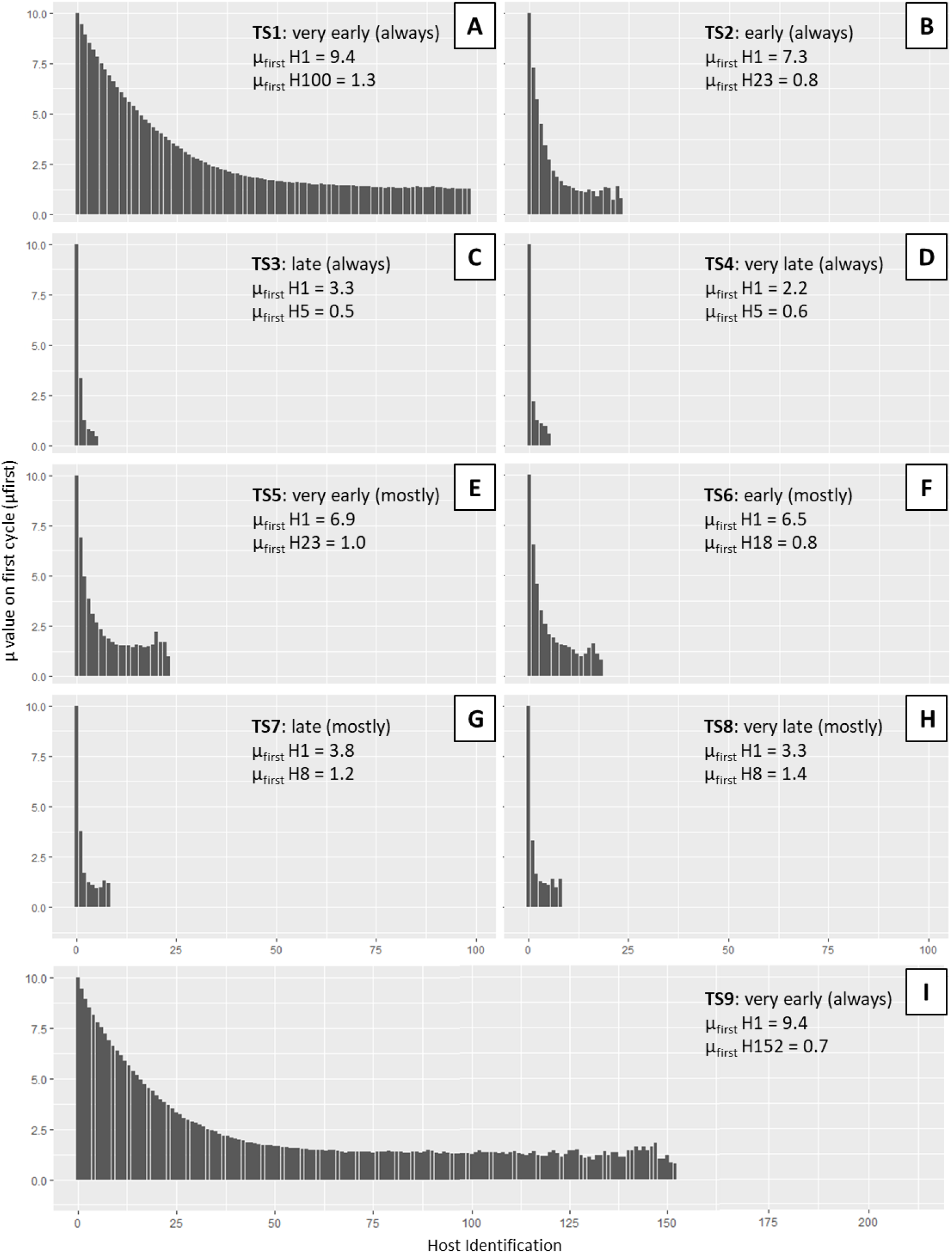
Mean µ values from the first replication cycle (µfirst) of the 1,000 simulations for each scenario, from TS1 to TS9. Each simulation (from TS1 to TS8) was programmed to reach 101 hosts and 44 replication cycles within each host by default in the transmission chain. The TS9 scenery was built with 350 simulations and 44 replication cycles for 501 hosts. The y axis, the “µ value on first cycle (µfirst)” label, shows the values of the replicative classes that ranges from zero to 10 and the x axis, “Host Identification”, indicates the amount of infected hosts in the simulations, ranging from the host zero (H0) to the host 100 (H100) and H152. The first host, H0, is the one that harbors the five higher fitness TF particles that replicate into a viral population and initiate the transmission chain. In the very early and early scenarios (TS1, TS2, TS5, TS6 and TS9) the following hosts still receive high fitness viral particles within a short transmission time. In the opposite way, for the late and very late transmission events (TS3, TS4, TS7 and TS8), the µ value suffers an abrupt decline in the second host already (H1). This indicates that the viral population within the donor host underwent several replication cycles and consequently several mutation events. For this reason, the TF particle features were lost and the high replication classes were diminished, reaching a shorter transmission chain. Each panel shows the µ values from the first replication cycle for the H1 and the last host infected in the summarized data text.

**Figure 6.**
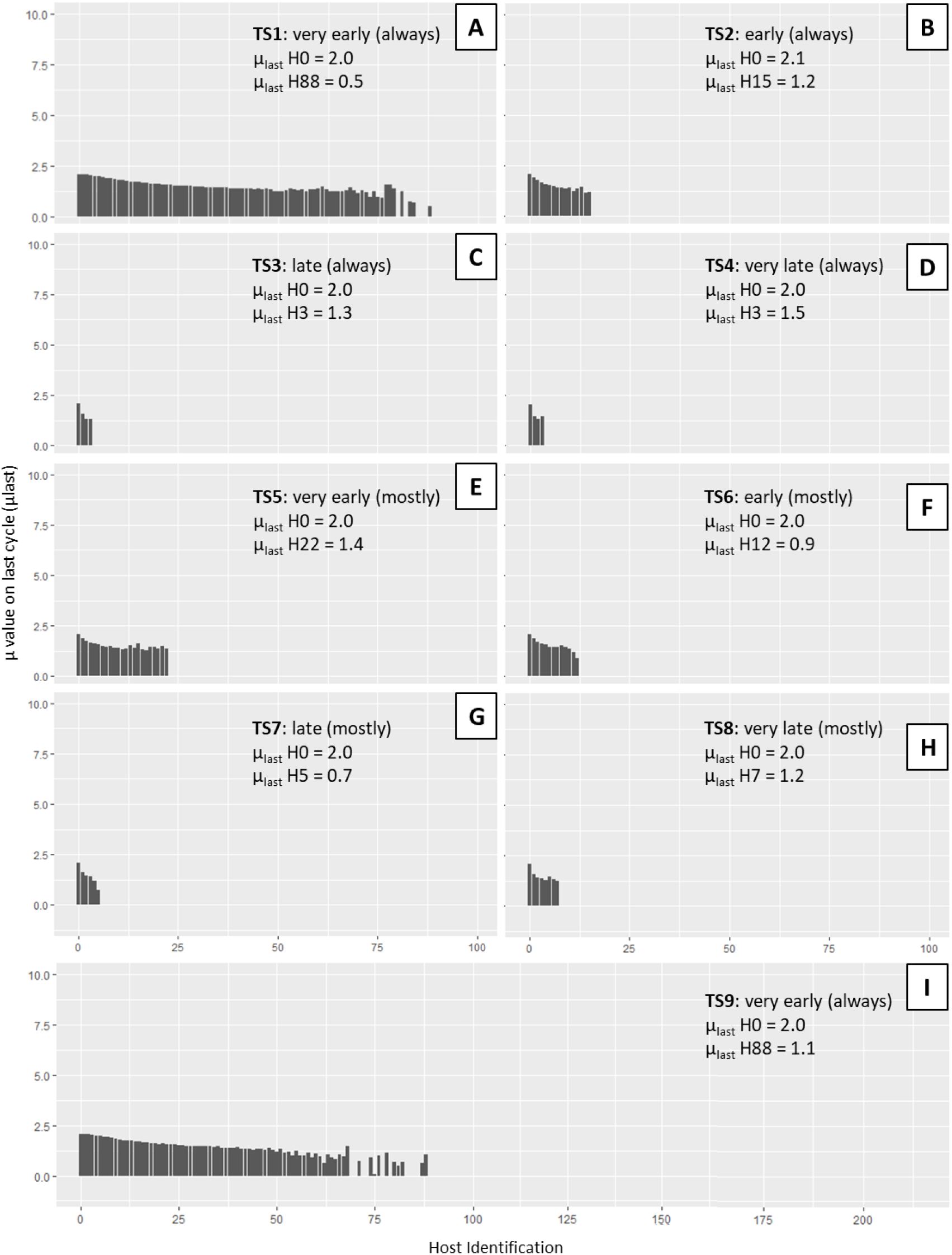
Mean µ values from the last replication (µlast) cycle (44th replication cycle) for the 1,000 simulations for each scenario, from TS1 to TS9. Each simulation (from TS1 to TS8) was programmed with 101 hosts and 44 replication cycles within each host by default in the transmission chain. The TS9 scenery was built with 350 simulations and 44 replication cycles for 501 hosts. The y axis, the “µ value on last cycle (µlast)” label, shows the values that range from zero to 10.0 and the x axis, “Host Identification”, indicates the amount of infected hosts in the simulations, ranging from the host zero (H0) to the host 100 (H100). The first host, H0, is the one that harbors the five higher fitness TF particles that replicate into a viral population and initiate the transmission chain. Note that the H0 already presents low fitness particles in the 44th replication cycle. This happens because at this final cycle the viral population is fairly adapted to the host environment and its selective pressures. It is worth mentioning that despite the transmission moment, at the 44th replication cycle, all hosts present similar viral particles with low fitness values. It emphasizes the importance of the very early/early timing for a successful TF transmission. Each panel shows the µ value for the H1 and the last host infected in the summarized data text.

It is important to notice that 1,000 simulations were performed for each TS, however not all simulations reached the last host in the transmission chain. For instance, in TS1 all simulations reached H20; 385 simulations reached H65; and only 84 simulations reached H100.

Scenario TS5 (very early transmission mostly) comprises very early transmission events, as well. Although, in this transmission scenario the transmission moments fluctuate among all ranges of replication cycles, but are mostly concentrated at the very early transmission times (cycles 0 to 4). The data generated by these simulations was similar to the simulation of TS1.

However the transmission chains ended earlier, reaching the maximum of 24 hosts (H0 to H23). The third transmission event at H3 presented Rmax=5.0 decreasing to Rmax=2.0 up to H23 (Fig. 4E). It was necessary only two transmission events for µ_first_ to attain the value 5.0 at H2 and H23 presented µ_first_=1.0 (Fig. 5E). In the last replication cycle, the branching process remained supercritical, with µ fluctuating from 2.0 to 1.4 (Fig. 6E).

Scenarios TS2 (early always) and TS6 (early mostly) displayed similar results, with a continuous decrease in the particle fitness and in the length of supercritical transmission chains. In TS2 the transmission chain displayed a slightly milder decrease of Rmax, with Rmax going below 5.0 at H4 and a also slightly longer transmission chain reaching H23 with Rmax=1.0 (Fig. 4B). While in TS6 Rmax became less than 5.0 at H3 and the supercritical transmission chain reached H18 with Rmax=1.0 (Fig. 4F). The same pattern was observed with the µ_first_ parameter, where TS2 kept µ_first_ above 1 up to H16, with µ_first_=1.2, and decreasing to µ_first_=0.8 at H23 (Fig. 5B). In both TS2 and TS6, the supercritical status was maintained till H12, with µ_first_=1.1, and decreasing to µ_first_=0.8 at H18 (Fig. 5F). The parameter µ_last_ of TS2 ranged from 2.1 to 1.2 over H0-H15, respectively (Fig. 6B). While, in TS6, µ_last_=2.0 at H0 and µ_last_=0.9 at H12 (Fig. 6F).

Scenario TS3 (late always) and TS7 (late mostly) exhibited similar results. Scenario TS3 reached H5 with Rmax=0.6 (Fig. 4C) and µ_first_=0.5 (Fig. 5C). For the parameter µ in the last replication cycle, it was possible to obtain data up to H3 only and µ varied from 2.0 to 1.3 (Fig. 6C). Scenario TS7 reached H8 with Rmax=2.0 (Fig. 4G) and µ_first_=1.2 (Fig. 5G). For the parameter µ in the last replication cycle, it was possible to obtain data until H5 only and µ varied from 2.0 to 0.7 (Fig. 6G).

Scenarios TS4 (very late transmission always) and TS8 (very late transmission mostly) exhibited similar results. Scenario TS4 reached H5 with Rmax=1.5 (Fig. 4D) and µ_first_=0.6 (Fig. 5D). The parameter µ_last_ ranged from 2 to 1.5 over H0 - H3, respectively (Fig. 6D). Scenario TS8 exhibited a slightly longer transmission chain reaching H9 with Rmax=0.0 (Fig. 4H) and with µ_first_=1.4 at H8 (Fig. 5H). The parameter µ_last_ ranged from 1.2 to 2.0 over H0 - H7 (Fig. 6H).

Scenario TS9 was defined with transmission events randomly occurring in the range of cycles 0 to 4 and transmission chains of length 501. Each host went through all the 45 replication cycles or all the and 350 simulations. The transmission chain was interrupted at H153. After H65, the parameter µ at the last replication cycle displayed oscillations from 2.0 to 0.0 (Fig. 5I).

## 4. DISCUSSION

It was possible to observe in scenario TS1 (very early transmission always) that the viral particles retained high enough fitness to successfully infect the total number of hosts in the simulations (101 hosts). The parameters Rmax and µ_first_ remained above 1 throughout the transmission chain (Fig.4A and Fig.5A, respectively) sustaining a supercritical status and allowing the population to maintain itself. The fitness of the viral population slowly decreases over time. That happened because the transmission cycles varied from 0-4, which was enough for the fitness to decrease a small amount per replication cycle through accumulation of deleterious effects.

In scenario TS5 (very early transmission mostly) it was possible to observe the negative impact on fitness that some late, or very late, transmissions provoked. Despite the fact that most transmissions occurred at very early moments, the deleterious accumulation effect of late and very late stages was enough to drastically reduce the length of transmission chains (Fig. 5E). Scenario TS5 reached a maximum number of 24 hosts in a chain (Fig. 4E). A transmission tree with 24 generations represents a population of 9.38×10^13^ individuals, according to equation (2).

Scenarios TS2 (early always) and TS6 (early mostly) showed similar results and analogous to scenarios TS1 and TS5, respectively. Clearly, TS2 and TS6 displayed smaller transmission chains in comparison to TS1 and TS5, since the prevalent transmission cycles went a little forward. Both scenarios were impaired by the deleterious effects from late transmission events despite their minor presence relative to early transmission events.

Scenarios TS3 (late always), TS7 (late mostly), TS4 (very late always) and TS8 (very late mostly) exhibited opposite effects in comparison to the very early/early “always”/”mostly” scenarios. More specifically, TS1 and TS2 exhibited a reduction in their transmission chain length when compared to TS5 and TS6, respectively. The observed effect was caused by the deleterious effect accumulated over the time. In the opposite direction, scenarios TS7 (Fig. 5G) and TS8 (Fig. 5H) exhibited slightly longer transmission chain length compared to TS3 (Fig. 5C) and TS4 (Fig.5D), respectively. For instance, TS8 reached H9, so the transmission tree represents a population of 350,000 individuals. The very early/early transmission events allowed TS7 and TS8 to explore more possibilities of the transmission tree, with TF particles of higher fitness. Both scenarios represent the consequences of the accumulation of deleterious effects. Late replication cycles allow a higher occurrence of deleterious mutations, thus the fitness of viral particles decreases over time. The fitness loss increases the occurrence of extinction events.

Scenario TS9 was an extra round of simulations of very early (always) transmissions events. Considering that TS1 simulations were set up to end at H100, it was unclear how much longer the transmission chains could be. The maximum length of transmission chains was 153. After that, the parameter µ became less than 1 and it was not possible to maintain the transmission of viable TF particles. The parameter Rmax of H153 was 0, which means that all 5 particles transmitted by H152 had Replicative Class R=0, thus ending the transmission chain. The parameter µ_last_ of each infected host in the transmission chain denotes the impact induced by the occurrence of deleterious mutations within each host (Fig. 6).

If at the last cycle the branching process is subcritical then any transmission event would transmit very low fitness particles to any new host. This is seen throughout all transmission scenarios: the parameter µ_last_ is bigger than 1 at H0 and decreases until a subcritical value along the transmission chain. The last host of the transmission chain carries the accumulated effect of deleterious mutations from all previous hosts in the chain. If the viral particles composing the initial inoculum display low fitness at the transmission bottleneck, it is likely that the viral population may become attenuated before being able to perform another infection.

Vrancken et al. [11] relate their findings to a continued transmission hypothesis, in which several hosts preserve the original transmitted founder particles or new high fitness transmitted founder particles, maintaining the circulation of the virus on the host population. This scenario implies very early and fast transmission events. The longer a TF particle remains in a host, the higher is the mutational pressure. Consequently, the original features of the TF particles are lost. This interpretation can be applied to the findings of Yu et al. [12], as well.

A genetic signature that characterizes TF particles has not yet been detected. Keele et al. [14] hypothesized that established infectious viral progenies may be identified in early stages as distinct genetic lineages presenting a low sequence diversity or that it may be possible to identify a consensus sequence from the early stage TF particles. The restricted amount of TF particles detected and the low genetic diversity present before the viremia peak could be interpreted as a finite window of vulnerability. That is, specific very early stage treatments could eliminate the TF particles, hence the viral population wouldn’t be able to establish itself in the new host.

After more than twenty years of research, it was found that people living with HIV under antiretroviral therapy with suppressed viral load are unable to transmit HIV via sexual intercourse. In 2018 UNAIDS proposed the term “undetectable = untransmittable” to refer to this situation [38]. The viral load is a consequence of the infection establishment and may indicate whether it was fully successful or not. It might be hypothesized that under these conditions, the TF particles are completely damaged and/or extinguished by the antiretroviral treatment. It is known that even under antiretroviral treatment, the infected individual’s viral load remains at a level of less than 50 copies/mL and it may suffer oscillations, known as “blips” [38–40]. Now, if one suppose that “blips” may occur and transmission events have vanishing probability in the “undetectable = untransmittable” situation, it is possible to consider that the transmitted viral particles generated during the blip events have already lost the TF features and are completely adapted to the current host. Therefore, only low fitness viral particles would be transmitted and they would not be able to establish a new successful infection. Accordingly, Crowell et al. [39], assume that the earlier the antiretroviral therapy starts, less blips the infected individual probably displays and vice-versa. Finally, in this regard, it is worth mentioning a few points on the sidelines of this discussion: (i) not all HIV infected patients present blips and (ii) the viral load oscillation is probably related to a reservoir effect, which is off-topic.

Also in this context, serodiscordant couples studies could provide similar insights. Gray et al. [41] followed a 174 serodiscordant monogamous couples cohort for 3 years and 11 months and samples/information were collected every 10 months. They verified that the transmission probability fluctuated proportionally to the detected viral load, with values of transmission probability of 0.0001/act and viral loads of 1,700 copies/ml or less; and 0.0023/act and 38,500 copies/ml or more. It is important to mention that the patients were not under retroviral treatment. Considering that the study did not provide precisely which stage the infection occurred in the donor patient, it is likely that the transmission probabilities for serodiscordant couples might be ruled by low fitness particles from late or very late stages of infection and not necessarily only act events. Recent studies of serodiscordant couples corroborate the “undetectable = untransmittable” status of people living with HIV, because the infected pair is usually under retroviral treatment and decline the use of condoms in a stable or exclusive sexual relationship [42–44].

Joseph et al. [8] described that there may be a high genetic variability in the donor host compared to the recently infected receptor host. This may favor the transmission event, increasing the probability of a TF particle to be in the transmitted bottleneck. This approach regarded the donor as a late stage host which presented a high particle diversity as a consequence of the infection moment. It is not considered that the viral particles may suffer a fitness decrease according to the mutation rates and the deleterious effects. To have a TF particle in late stages requires a mechanism that protects them from mutations. Latent reservoirs could give this protection to TF particles for some time [45]. Nevertheless, matching the release of TF particles from the reservoir with the transmission bottleneck seems to be a very low probability event. Boeras et al. [19] described that there may exist a process that targets the best fitness particles and leads them to transmission.

Several research groups discuss the mode and timing of HIV transmission and researchers in the field disagree about the most relevant transmission moment for the success of viral infection [6, 7, 11, 12, 46–49]. There is a broad range of time intervals that can be considered as early or late stages. Given that the replication cycle of the HIV is 2.6 days, an interval of months presents a large amount of mutation events, for instance. Furthermore, none of the above-cited papers assign a fitness feature to the transmitted founder particles. Time is usually considered only as a transmission moment, yet irrelevant for the replication cycle process and the consequent mutation events are neglected, as well. Only the contact rates and transmission stages are considered as the main variables. The point is not that the amount of variables considered is not appropriate, but about the absence of the fitness of the particles as one of the main variables to be considered.

It is important to emphasize that this work demonstrates a possible induced impact on the fitness of transmitted viral particles through the analysis of a simple model. The low fitness viral particles present in the last replication cycle of each host does not imply that there is a natural viral infection attenuation. The fitness observed in the data generated reflects a “measure” of how probable a new transmission event is to occur and how likely it is for the infection to establish itself or not. This observation is restricted only to the transmission and infection establishment potential. It is not a fitness index for the not transmitted viral population.

Finally, we mention some important aspects of viral evolution and epidemiology that are not taken into account by out model: (i) latent reservoirs, (ii) defective-interferent particles and their possible complementation mechanisms, (iii) recombination of viral particles, (iv) pathogen to host adaptation leading to increase of infectivity (this is important for zoonotic diseases), (v) antiretroviral therapy intervention. Of course, it is possible to extend the model and the program to include some of these features. We will leave the investigation of such extensions for other publications.

## 5. CONCLUSIONS

Mathematical and computational models have been widely applied to epidemiological studies to shed light on its mechanisms, to make predictions about the range of infections in host populations and to develop vaccine strategies, for example [50–53]. HIV transmission routes are well known, however, studying the exact instant of the transmission event presents several limitations. The access to biological samples in the precise moment of the viral population establishment and development and/or the size of the studied host population are very strong limitations. Accordingly, the *in silico* model developed in this paper is a tool for the analysis of transmission events in a host population using data from the dynamics of intra-host virus populations. The ENV-TF program enables a plethora of simulation scenarios by variation of its user defined parameters. It is possible to isolate the effect of certain variables and study the impact of each of them at a time. The simulation sample sizes, both of the host population and the viral populations in each host, is only limited by the computer processing power available.

The simulation results show the impact of fitness loss in the viral population due to the accumulation of deleterious effects through transmission events among hosts. The very early transmissions indicate that there may exist an optimal moment for the viral transmission to occur.

For instance, in the case of HIV, this is a critical moment, because there may be no suspicion about an infection due to absence of clinical symptoms. In fact, clinical symptoms of HIV infection only manifest within 3-6 weeks and can be very similar to flu symptoms [35, 54]. Hence, the diagnostics may be postponed, implying an untrue health status and a possible sexual risk behavior.

As discussed before, our simulated data are in agreement with several findings in the literature. Early infections probably imply a lower fitnes loss and maintenance of TF characteristics for a longer time, allowing a higher number of successful infection events. The short optimal transmission period of early stages may be a meaningful process that could sustain a viral epidemic or pandemic. In a general way, the mechanism based on TF particles does not support that chronic stage infections are the main responsible for the maintenance of the long term circulation of the disease. In the chronic stage a high viral load may remain for a long time within the host. However, it is not clear if, in the chronic stage, there would exist functional TF particles and how they are selected to establish an infection in a new host.

## ACKNOWLEDGMENTS

We thank Diogo Castro dos Santos for the development of the ENVELOPE program and Marcos Saito de Paula for his contributions to the development of the ENV-TF code.

## AUTHOR CONTRIBUTIONS

Conceived and designed the experiments: LMRJ and FMA.

Performed the experiments: TNF and FMA.

Analyzed the data: TNF, FMA, IMVGC and LMRJ.

Wrote the paper: TNF, FMA, IMVGC and LMRJ.

## CODE AVAILABILITY

The complete code is available on github: https://github.com/antonelijr/ENV-TF

